# Endocytosis Inhibitors Block SARS-CoV-2 Pseudoparticle Infection of Mink Lung Epithelium

**DOI:** 10.1101/2023.07.12.548725

**Authors:** Ann Song, Rattapol Phandthong, Prue Talbot

**Author notes:** **Correspondence:** Prue Talbot.

## Abstract

Both spill over and spill back of SARS-CoV-2 virus have been reported on mink farms in Europe and the United States. Zoonosis is a public health concern as dangerous mutated forms of the virus could be introduced into the human population through spillback. The purpose of our study was to determine the SARS-CoV-2 entry mechanism using mink lung epithelial cell line (Mv1Lu) and to block entry with drug inhibitors. Mv1Lu cells were susceptible to SARS-CoV-2 viral pseudoparticle infection, validating them as a suitable disease model for COVID-19. Inhibitors of TMPRSS2 and of endocytosis, two pathways of viral entry, were tested to identify those that blocked infection. Dyngo4a, a small molecule endocytosis inhibitor, significantly reduced infection, while TMPRSS2 inhibitors had minimal impact, supporting the conclusion that the entry of the SARS-CoV-2 virus into Mv1Lu cells occurs primarily through endocytosis. The small molecule inhibitors that were effective in this study could potentially be used therapeutically to prevent SARS-CoV-2 infection in mink populations. This study will facilitate the development of therapeutics to prevent zoonotic transmission of SARS-CoV-2 variants to other animals, including humans.

## 1 Introduction

The emergence of COVID-19, which is caused by the SARS-CoV-2 virus, has posed a global health emergency (Hoffman et al. 2020; Li et al. 2020). As of now, SARS-CoV-2 virus has infected over 103 million people and caused more than 1.1 million deaths in the U.S. (cdc.gov and WHO). 69% of the U.S. population has been vaccinated, which has improved health outcomes and decreased the mortality rate. Nevertheless, COVID-19 is still a major health concern that kills about 200 - 400 people in the U.S. weekly, reoccurs in vaccinated individuals, and mutates rapidly producing new variants (Vignier et al. 2021). The highly infectious SARS-CoV-2 virus can enter human host cells via two pathways (Figure 1) (Hoffman et al. 2020; Cai et al. 2020). In the protease-based pathway, the receptor binding domain (RBD) in the viral spike (S) protein binds to ACE2 on the host cell, followed by cleavage of the S protein by TMPRSS2 and subsequent fusion of the virus and host cell. The second pathway involves endocytosis of the virus following the binding of the S protein to ACE2 (Bayati et al. 2021).

**Figure 1.**
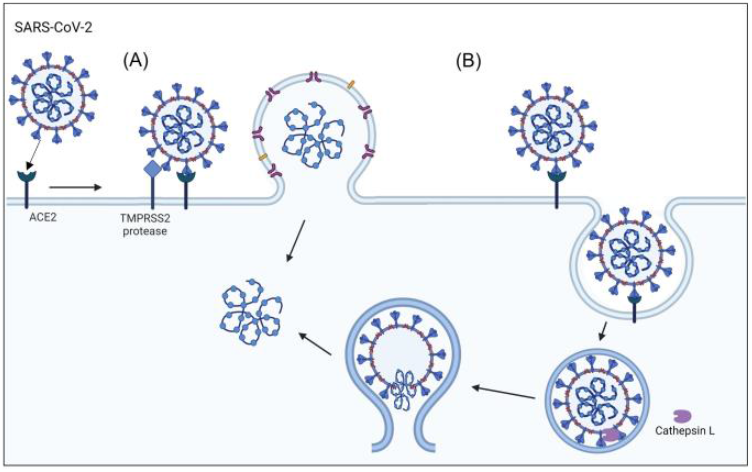
SARS-CoV-2 viral entry mechanisms. The SARS-CoV-2 virus binds to the ACE2 host cell receptor then either (A) TMPRSS2 processing of the viral spike protein leads to the fusion of the virus to the cell membrane or (B) the virus is taken up by endocytosis.

While the transmission of SARS-CoV-2 virus has been mainly among humans through aerosol (Zhou et al. 2020), zoonotic transmission of the virus can also occur. For example, SARS-CoV-2 transmission has been reported in wild animals (deer, hippo), livestock (mink), and house pets (cat, dog, hamster) (Shi et al. 2020; Zhao et al. 2020; Palmer et al. 2021; Hale et al. 2021). Reservoirs of virus in non-human species is a concern since new variants, which may be vaccine resistant, can develop and spill back to humans (Koopmans 2021). While most COVID-19 research has been on humans, the importance of zoonotic transmission has recently been appreciated by the scientific and public health communities (Madhusoodanan 2022). To prevent zoonotic transmission of the virus, it is necessary to understand the mechanism of viral entry into cells and identify drugs that inhibit infection in zoonotic species.

SARS-CoV-2 transmission in mink (*Neovison vison*) is the focus of this study. Mink, which are part of the Mustelidae family with weasels and ferrets, are among the mammals that are readily infected with SARS-CoV-2 virus (Reviewed in Salajegheh Tazerji et al. 2020). SARS-CoV-2, which is a species of the Coronoviridae family, includes viruses that cause SARS and MERS, which have known zoonotic origins (Wong et al. 2020). The first SARS-CoV-2 infection in mink was reported in the Netherlands (Molenaar et al. 2020), followed by Denmark, Spain, Italy, Sweden, and Greece (Boklund et al. 2021). The USDA also reported positive cases in mink in Utah (Cahan 2020). SARS-CoV-2 virus can replicate efficiently in the upper and lower respiratory tracks of mink, displaying symptoms similar to those of humans, such as interstitial pneumonia with alveolar damage (Molenaar et al. 2020). Since mink are farmed on a large scale for their fur, there is growing public health concern about variant transmission back to humans (Oude Munnink et al. 2021). Consequently, many farms have shut down, and the Dutch government banned mink farming in 2021. Although the origin of SARS-CoV-2 virus is unknown, Chinese fur factories, which produce half of the world’s pelts from live animal farms including mink, are considered a possible site of origin (Cohen 2021). Early in the pandemic, culling was the only way to stop mink zoonotic transmission and protect public health. Nevertheless, mink are still farmed in five countries, and the potential for zoonotic transmission from mink to other animals and humans still exists.

Spillback of COVID-19 from mink to human has been reported (Hammer et al. 2021; Lu et al. 2021). Although there is no evidence that Cluster 5 (mink-associated variant) spillback was associated with increased severity or transmissibility (Lassauniere et al. 2021), mink spillback should not be underestimated since future variants could be more dangerous. In a laboratory experiment, mink were permissive to the Omicron variant (BA.1), which is known for its rapid transmission and replication (Virtanen et al. 2022). The USDA has authorized testing an animal vaccine on some mink farms to establish data on the efficacy and safety that are required for approval. Vaccination provides active immunity by enabling antibody production, which reduces the severity of the disease and the chance of transmission. However, vaccination does not prevent infection, which could be blocked by drugs that target viral entry mechanisms.

The 3D structure and binding affinity of SARS-CoV-2 virus to ACE2 host receptors have been used to predict the susceptibility to infection in non-human animals (Damas et al. 2020; Sun et al. 2021; Hayashi et al. 2020). However, there are limitations to *in silico* studies that must be validated by experimental and clinical work. Damas et al. (2020) predicted the susceptibility of mink to SARS-CoV-2 virus to be very low based on structural analyses. Yet, infection of mink has been reported worldwide in mink farms, where animals live at high densities, a factor that may not have been considered in computational studies (Molenaar et al. 2020; Hayashi et al. 2020; Cahan 2020). The limitations of computational studies and the absence of cellular data on SARS-CoV-2 infection in mink are major gaps in our knowledge.

The goal of this study was to determine the mechanisms by which mink become infected with the human SARS-CoV-2 virus. Specifically, we determined the SARS-CoV-2 viral entry mechanism in mink cells and identified drugs that prevent infection. Our goals were accomplished using a SARS-CoV-2 lentiviral-based pseudotyping assay to assess viral entry into mink lung epithelial cells (Mv1Lu). Small molecule inhibitors targeting TMPRSS2 or endocytosis, the main viral entry pathways, were evaluated for their efficacy. The results are among the first to examine the SARS-CoV-2 entry mechanism in a non-human species and may inform public health policy.

## 2 Materials and methods

### 2.1 ACE2 Sequence Alignment

The amino acid sequences of ACE2 from selected species were obtained from the ENSEMBL (mink) and UniProt (other species) databases and aligned using NCBI blastp. The overall sequence identity and the residues involved in the ACE2-virus binding interface were compared with the human sequence.

### 2.2 Tissue Culture and Reagents

A fetal mink lung type II alveolar epithelial cell line, Mv1Lu, (ATCC, Manassas, VA) was grown in Eagle’s Minimum Essential Medium (EMEM) supplemented with 10% fetal bovine serum (FBS; Gibco, Carlsbad, CA) and 1% penicillin-streptomycin. HEK 293T and its ACE2-overexpressing cell line (HEK 293T-ACE2) (ATCC, Manassas, VA) were grown in DMEM with high glucose and 10% FBS. Both cell types were grown in T25 flasks until they reached approximately 90% confluency. For passaging, cells were washed once with phosphate-buffered saline (PBS), removed from the flask with 1.25 mL of 0.25% trypsin-EDTA for 3 min at 37 °C, and then centrifuged to remove the supernatant. The pellet was resuspended in complete growth medium.

The following small molecule inhibitors were used: ambroxol (10 µM; TCI Chemicals, Portland, OR), aprotinin (10 µM; Tocris, UK), Camostat (10 µM; Sigma-Aldrich, Burlington, MA), Nafamostat (10 µM; TCI Chemicals, Portland, OR), Dyngo 4a (20 µM, Selleckchem, Houston, TX), Chlorpromazine (CPZ) (10 µM; Sigma-Aldrich, Burlington, MA), Pitstop2 (20 µM; Abcam, Cambridge, UK), mβCD (20 µM; Sigma-Aldrich, Burlington, MA), genistein (20 µM; TCI Chemicals, Portland, OR), nystatin (20 µM; Sigma-Aldrich, Burlington, MA), and Filipin (20 µM; Sigma-Aldrich, Burlington, MA).

### 2.3 Protease Activity Assay

Fluorogenic substrates were prepared using appropriate reaction buffers. For furin cleavage, Pyr-Arg-Thr-Lys-Arg-AMC (1 mM, Bachem) was prepared in 100 mM HEPES buffer (pH 7.0), 0.5% Triton X-100, 1mM calcium chloride, and 1mM β-mercaptoethanol. For cathepsin L cleavage, Z-Phe-Arg-AMC HCl (1 mM, Bachem) was prepared in 50 mM Tris (pH 8) and 150 mM NaCl. For TMPRSS2 cleavage, Boc-Gln-Ala-Arg-AMC HCl (2.5 mM, Bachem) was prepared in 50 mM Tris (pH 8) and 150 mM NaCl.

For furin and cathepsin L assays, whole cell lysates were used. Mv1Lu cells were collected at a density of 1 × 10^7^ cells, washed twice with PBS, and lysed for 1 min on ice in RIPA buffer. The cells were then sheared with a 21-gauge needle, followed by centrifugation at 3,000 rpm for 5 min at 4 °C. The supernatants (4 µL) were added to each reaction well.

For TMPRSS2, live cells were used. Mv1Lu cells were seeded at a density of 2.5 × 10^4^ cells in a 96-well plate. Cells were preincubated with TMPRSS2 inhibitors for 24 h before measuring enzymatic activity.

The fluorogenic substrate was added to each well (final concentration: 100 µM for furin and cathepsin L, 10 µM for TMPRSS2). Fluorescence intensity was measured at 340/440 nm using a BioTek Synergy HTX, multi-mode microplate reader (Winooski, VT).

### 2.4 Immunofluorescence Microscopy

The cells were seeded in 8-well chamber slides (Ibidi; Grafelfing, Germany). When they reached confluence, the cells were fixed at room temperature in 4% paraformaldehyde (PFA) in PBS for 15 min, washed with PBS, and permeabilized with 0.1% Triton X-100 for 10 min. The cells were then blocked with 10% donkey serum. Primary antibodies were diluted in blocking buffer. The following primary antibodies were used: goat anti-ACE2 (1:200; R&D Systems, Minneapolis, MN), mouse anti-TMPRSS2 (1:200; Santa Cruz Biotechnology, Dallas, TX), and mouse-EEA1 (1:200; BD Biosciences, San Jose, CA).

Following overnight incubation with primary antibody at 4 °C, the cells were washed three times with 0.2% Tween in PBS. Next, the cells were incubated in the dark with Alexa Fluor-conjugated fluorescent secondary antibodies (1:500; Invitrogen, Carlsbad, CA). After washing in PBS, the cells were mounted with diluted Vectashield and DAPI for nuclear staining. A Nikon Eclipse Ti microscope was used to image at 60x in water immersion. The NIS Elements Software and ImageJ software were used to image and analyze the data.

### 2.5 RNA Extraction and RT-PCR assays

RNA was extracted from Mv1Lu cells using the RNeasy Mini Kit (Qiagen, Germantown, MD) according to the manufacturer’s protocol. Primer sequences from Heller et al. (2006) are provided below:

hACE2 F: 5’- CTTGGTGATATGTGGGGTAGA -3’

hACE2 R: 5’- CGCTTCATCTCCCACCACTT -3’

mink ACE2 F: 5’- GGAACTCTACCATTTACTTACA -3’

mink ACE2 R: 5’- TCCAAGAGCTGATTTTAGGCTTAT -3’

hGAPDH F: 5’- CATCACCATCTTCCAGGAGC -3’

hGAPDH R: 5’- CTTACTCCTTGGAGGCCATG -3’

RT-PCR was performed after generating first-strand cDNAs from 2 µg of total RNA using iScript Reverse Transcription Supermix (Bio-Rad, Hercules, CA), as recommended by the manufacturer. Diluted cDNA (40 ng/µL) was amplified using Phusion Polymerase (Thermo Fisher Scientific, Waltham, MA). The samples were analyzed using agarose gel electrophoresis (1%).

### 2.6 Lentiviral Production to Generate SARS-CoV-2 Pseudoparticles

HEK293T cells were plated with antibiotic-free medium at a density of 7 × 10^6^ cells in a T75 flask and transfected via Lipofectamine3000 (Invitrogen, Carlsbad, CA) using a total of 15 µg of BEI lentiviral plasmids (Table 1), 60 µL of Lipofectamine, and 20 µL of P3000 per T75 flask following the manufacturer’s protocol. After overnight incubation, fresh medium was added to cells supplemented with 1% BSA. Fluorescence microscopy was used to observe the transfection efficiency based on Zsgreen expression.

**Table 1.**
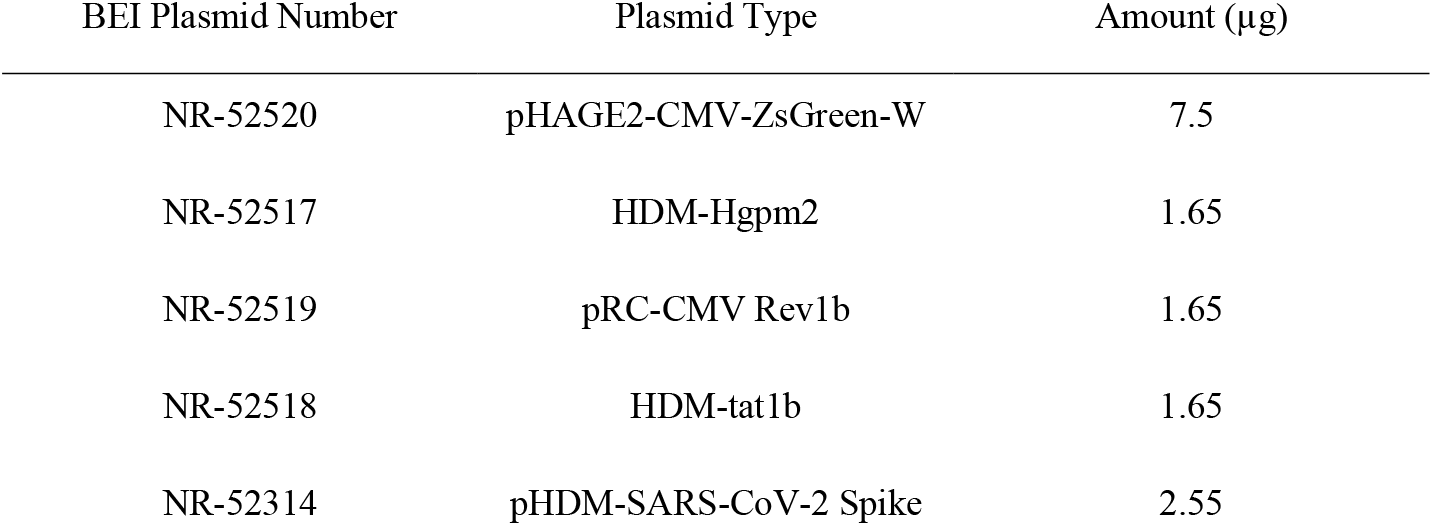
Summary of BEI Lentiviral Vectors Used For Transfection

Conditioned medium was collected 48 h post-transfection. The conditioned medium was centrifuged, and the supernatant was filtered using a 0.45 µm syringe filter. The filtered supernatant was mixed with 5x PEG (Abcam, Cambridge, UK) and precipitated overnight at 4°C. The lentivirus was collected by centrifugation and resuspending the pellet in Viral Re-suspension Solution (Abcam, Cambridge, UK). Virus aliquots were stored at -80 °C. Prior to the experiments, the transduction efficiency of each batch of viruses was tested via flow cytometry of infected HEK 293T-ACE2. Viral titers were calculated using the Poisson formula (Crawford et al. 2020; Song et al. 2023). The flow cytometry conditions were the same as those for the Mv1Lu cell protocol. The medium was collected from wells containing cells that were not transfected, and PEG-precipitated as described above, and then used for mock infection.

### 2.7 Pseudotyping of human SARS-CoV-2 viral pseudoparticles

Mv1Lu cells were plated at a density of 30% and infected with SARS-CoV-2 viral pseudoparticles (multiplicity of infection, or MOI, of 0.1). The medium was replaced after overnight incubation of the infected cells. Fluorescence microscopy was used to observe transduction efficiency. Cells were trypsinized 48 h post-infection and washed three times with 0.5% BSA in PBS. After the final wash, the cells were resuspended in the same wash buffer and analyzed using flow cytometry.

Novocyte flow cytometer was used to detect ZsGreen in the FITC channel. The resulting flow cytometer files were analyzed using NovoExpress software. Mock infection was used as a background control. Prior to infection, the cells were preincubated for 2 h with TMPRSS2 inhibitors or endocytosis inhibitors, which were kept in the medium until harvesting for flow cytometry.

### 2.8 Endocytosis Assay

Mv1Lu cells were seeded in 8-well chamber slides. The cells were incubated overnight in Dyngo4a and TRITC-conjugated Dextran (Invitrogen, Carlsbad, CA). After fixation in 4% PFA, mounting was performed using diluted Vectashield with DAPI. Immunocytochemistry with an antibody against EEA1 was also performed to further validate dextran uptake, as described above for ACE2 and TMPRSS2.

### 2.9 Data Analysis and Statistics

The means and standard errors of the mean for three biological experiments were plotted using GraphPad Prism 7 software (GraphPad, San Diego, CA). For infection data, the mean of the DMSO group was set to a value of 100, and the inhibitor groups were compared to this value. Statistical significance was determined using Minitab Statistics Software (Minitab, State College, PA). When the data did not satisfy the assumptions of ANOVA (normal distribution and homogeneity of variances), they were subjected to a Box-Cox or logarithmic transformation. All infection analyses were performed using a one-way ANOVA, and TMPRSS2 cleavage analysis was performed using two-way ANOVA, in which the factors were time and treatment. When the ANOVA means were significant (p<0.05), groups treated with inhibitors were compared to the DMSO control group using Dunnett’s post hoc test.

## 3 Results

### 3.1 ACE2 in mink and human have similar amino acid sequences

15 out of 16 mink ACE2 amino acid orthologs had more than 80% sequence similarity with human ACE2 (Table 2). Mink showed a high sequence identity (82.99%), like other known susceptible species, such as cats and dogs (Shi et al. 2020). Since ACE2 residues that directly bind to the RBD in the S protein have been identified (Lan et al. 2020), these residues in ACE2 were aligned in various vertebrates to predict their affinity to the viral S protein (Figure 2). Lysine residues 31 and 353 are particularly critical for binding to the receptor binding domain of the S protein (Wan et al. 2020). In mink and other mammalian species that are readily infected by SARS-CoV-2, these critical residues were identical to those in humans. However, in species that are less susceptible, residues 31 (mouse, chicken, and bat) and 353 (mouse and rat) were different from human ACE2.

**Table 2.** ACE2 is highly conserved in various mammalian species.

| Animal | % Identity |
| --- | --- |
| Cat | 85.22 |
| Rabbit | 85.14 |
| Hamster | 84.26 |
| Dog | 83.50 |
| **Mink** | **82.99** |
| Deer | 82.44 |
| Rat | 82.37 |
| Hippo | 81.86 |
| Mouse | 81.86 |
| Sheep | 81.74 |
| Goat | 81.74 |
| Bovine<sup>1</sup> | 81.48 |
| Pig | 81.37 |
| Cattle<sup>2</sup> | 80.99 |
| Bat | 80.73 |
| Chicken | 65.97 |

<sup>1</sup> Hybrid of *Bos indicus* and *Bos taurus*
<sup>2</sup> *Bos taurus*

**Figure 2.**
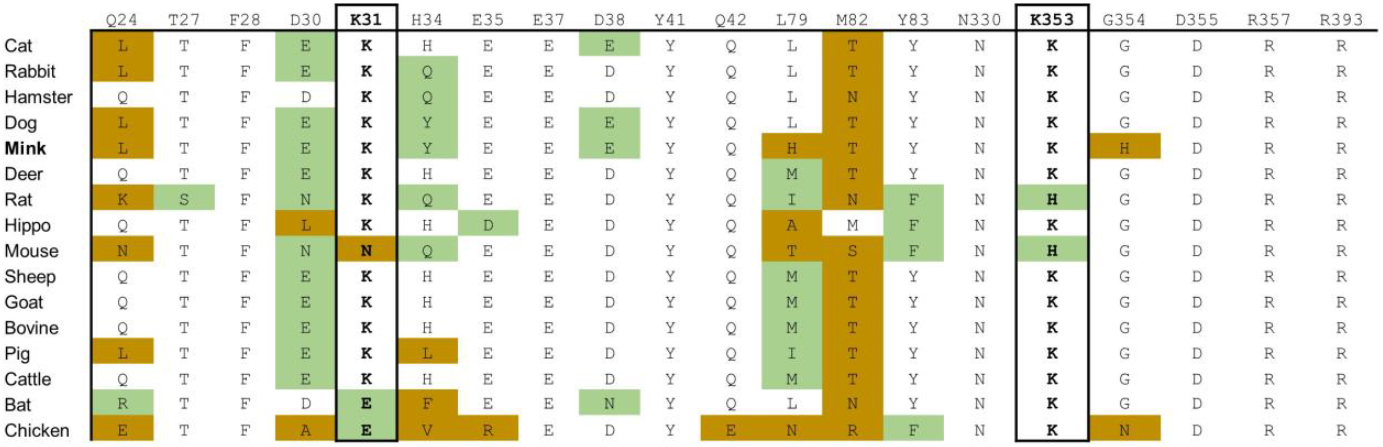
Cross-species conservation of ACE2 at known binding residues. The ACE2 amino acid sequences of different vertebrate species were aligned, and the amino acids found at each of the residue position involved in binding are shown. Amino acid substitutions are colored as either non-conservative (orange) or conservative (green) based on their similarity to the human residue.

### 3.2 Mv1Lu cells express the machinery for SARS-CoV-2 infection

To support high conservation and verify expression, the ACE2 receptor was examined in mink Mv1Lu cells using RT-PCR and immunocytochemistry. RT-PCR was performed using a human ACE2 primer set (hACE2) and mink ACE2 primer set (MvACE2) to amplify the ACE2 transcript (Figure 3A). The HEK 293T-ACE2 and HEK 293T cell lines (primer controls) expressed the ACE2 gene, verifying the RT-PCR assay. Mink cDNA reverse-transcribed from Mv1Lu RNA showed robust amplicons for both primer sets, indicating the presence of the ACE2 transcript. Mv1Lu cells labeled with the ACE2 antibody had numerous puncta over their surfaces, demonstrating translation of the ACE2 transcript (Figure 3B). Cells labeled with donkey anti-goat secondary antibody alone (control) had no fluorescent labels (Figure 3C).

**Figure 3.**
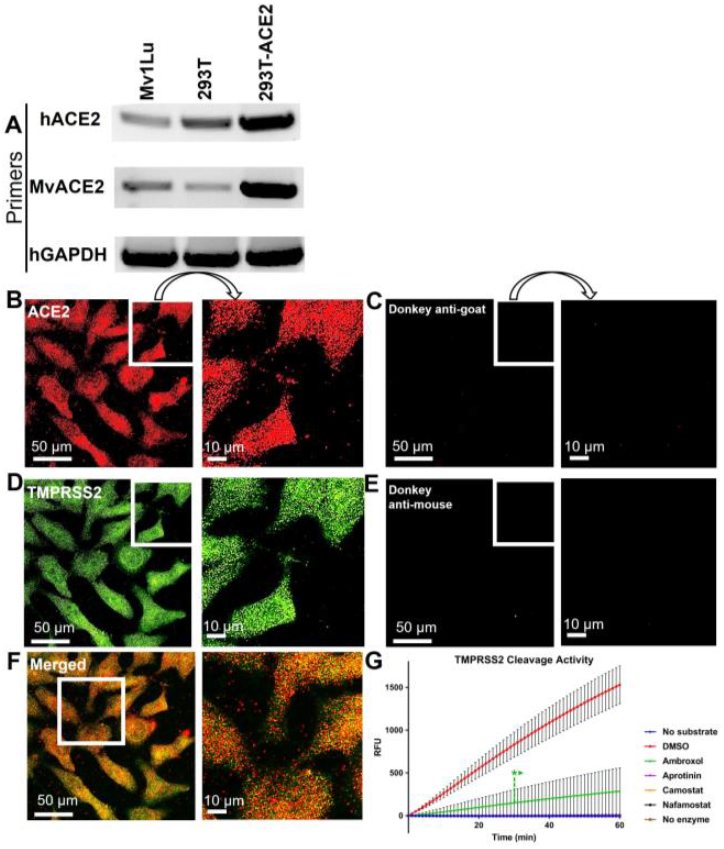
Mv1Lu cells are susceptible to SARS-CoV-2 pseudoparticle infection. (A) Representative RT-PCR analysis of ACE2 expression in Mv1Lu cells and HEK293T cells, showing primers in the rows and cell lysates in the columns. hACE2 = human ACE2 primers; MvACE2 primers = mink ACE2 primers; hGAPDH = housekeeping control; 293T = RNA extracted from HEK 293T; 293T-ACE2 = RNA from HEK 293T cells stably overexpressing ACE2. Mv1Lu cells expressed the ACE2 transcript using both primer sets. (B -F) Immunocytochemistry showing that Mv1Lu cells expressed ACE2 (B, C) and TMPRSS2 (D, E). Three independent experiments were performed. (G) TMPRSS2 activity in the presence of small molecule inhibitors. Mv1Lu cells had high TMPRSS2 activity, which was also lowered with inhibitors. DMSO = solvent control; RFU = relative fluorescence unit or normalized fluorescence intensity. In G, each group is the mean ± SEM.

The TMPRSS2 protease was expressed in Mv1Lu cells and appeared as numerous puncta on the surface of immunolabeled cells (Figure 3D). Cells labeled with secondary donkey anti-mouse antibody-only control did not show fluorescent puncta (Figure 3E). Many of the ACE2 and TMPRSS2 puncta were colocalized in the Mv1Lu cells (Figure 3F).

The activity of the proteases involved in SARS-CoV-2 infection [furin, cathepsin L (Supplementary Figure 1) and TMPRSS2 (Figure 3G)] was detected in the lysates or live cells as indicated by the strong signal emitted from the fluorogenic substrates. While all three enzymes showed high activity, we only followed up with TMPRSS2 since it is involved in the initial infection step, unlike furin and cathepsin, which function in post-viral entry processes.

Three TMPRSS2 inhibitors (aprotinin, Camostat, and Nafamostat) completely reduced TMPRSS2 activity, whereas ambroxol reduced activity to 20% of the DMSO control (Figure 3G). These data show that these inhibitors were effective in blocking TMPRSS2 activity in Mv1Lu cells.

### 3.3 SARS-CoV-2 enters Mv1Lu cells via endocytosis

Human SARS-CoV-2 pseudoparticles were produced using previously published procedures (Crawford et al. 2020; Song et al. 2023). Generated pseudoparticles were first titered with HEK293T-ACE2 cells using flow cytometry to detect the ZsGreen viral backbone (Figure 4A-C). High viral titers were obtained using 0.625 µL of pseudoparticles, resulting in a transduction efficiency of 53.44% (Figure 4A). 2-fold serial dilutions of the pseudoparticles showed a gradual decrease in transduction: 29.99% with 0.3125 µL and 15.55% with 0.15625 µL (Figure 4B-C). The MOI was calculated using the Poisson equation to assess the SARS-CoV-2 pseudoparticle infection in Mv1Lu cells (Crawford et al. 2020). Viral infection in Mv1Lu cells showed 67.49% transduction efficiency, with an MOI of 0.3 (Figure 4D-E). The MOI was adjusted to 0.1 for all other infection assays. These results indicated that Mv1Lu cells are highly susceptible to human SARS-CoV-2 pseudoparticle infection.

**Figure 4.**
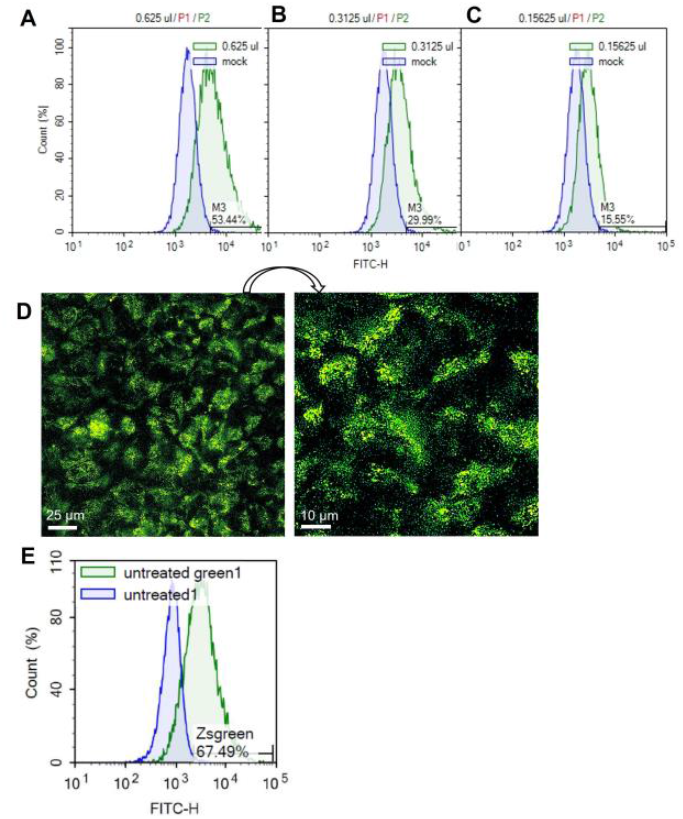
SARS-CoV-2 pseudoparticle infection in Mv1Lu cells. (A-C) Flow cytometry was performed to calculate the MOI using two-fold serial dilutions of the pseudoparticles in HEK 293T-ACE2 cells. Histograms of total cell population (Count %) vs Zsgreen signal (FITC-H) were plotted. M3 indicates the percentage of Zsgreen-positive cells compared to the mock control in each sample. (D) Representative fluorescent image of infected Mv1Lu cells. Mv1Lu cells were Zsgreen-positive, indicating infection with SARS-CoV-2 pseudoparticles. (E) Flow cytometry showed that 67.49% of Mv1Lu cells were infected compared to the untreated control.

TMPRSS2 inhibitors were used to determine if they could reduce infection in Mv1Lu cells (Figure 5A). The mean infection in DMSO was set to 100, and the inhibitor groups were compared to this value. Statistical analysis showed that none of the protease inhibitor groups were significantly different from the DMSO control.

**Figure 5.**
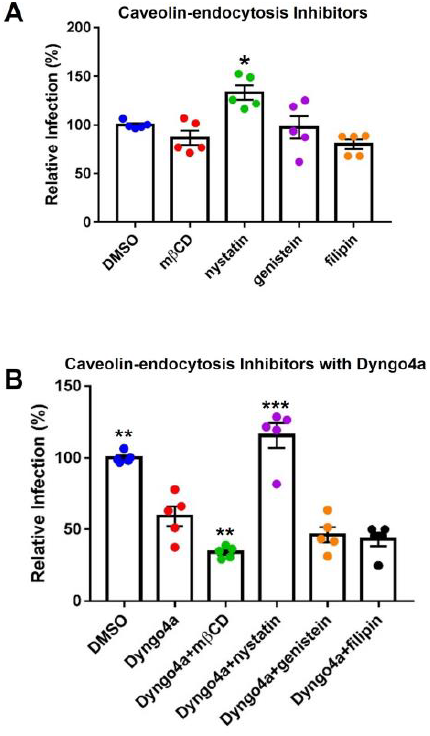
Effects of inhibitors on Mv1Lu infection using SARS-CoV-2 pseudoparticles. Data are normalized to the DMSO control group. (A) SARS-CoV-2 pseudoparticle infection was not significantly affected by TMPRSS2 inhibitors. (B-E) Activity of inhibitors against endocytosis was assessed using Dextran (TRITC conjugated) or (F-I) the EEA1 antibody. Endocytosis inhibitors lowered endosome formation. (J) An endocytosis inhibitor, Dyngo4a, significantly lowered infection. Pitstop2 and CPZ were not significantly different than DMSO. Data are the means ± SEM of three independent experiments. *p<0.05. One-way ANOVAs were performed on raw data.

To test the possibility of viral entry via endocytosis, Mv1Lu cells were treated with three endocytosis inhibitors (Dyngo4a, Pitstop2, and CPZ). Endocytosis was observed using dextran conjugated to TRITC, which fluoresces red upon uptake by endocytosis. In the DMSO control group, dextran-TRITC was taken up by the cells and appeared as small fluorescent puncta, consistent with its uptake by endocytosis (Figure 5B). Cells treated with endocytosis inhibitors (Dyngo4a, Pitstop2, and CPZ) showed a significant decrease in the uptake of dextran-TRITC (Figure 5C-E). Cells treated with Dyngo4a or CPZ showed little to no fluorescence, whereas those treated with Pitstop2 showed some evidence of endocytosis.

The EEA1 antibody was used to track early endosome formation. Endosomes were detected as small green puncta in the DMSO control group (Figure 5F). When cells were treated with Dyngo4a, Pitstop2, or CPZ, endosome formation was significantly reduced (Figure 5G-I). While endosomes were observed in the CPZ group, their numbers were far lower than in the DMSO control group.

Endocytosis inhibitors were next used to determine if they could reduce infection relative to the DMSO control (Figure 5J). Dyngo4a and Pitstop2 both reduced infection relative to the DMSO control, but only Dyngo4a was significant (p = 0.028). CPZ increased infection but was not significantly different from the control.

### 3.4 SARS-CoV-2 pseudoparticles infected Mv1Lu cells via endocytosis

The TMPRSS2 and endocytosis pathways were simultaneously targeted using cocktails of inhibitors. When statistical significance was found, Dunnett’s post hoc test was used to compare the groups to Dyngo4a alone. As seen before (Figure 5J), Dyngo4a alone significantly lowered infection compared with DMSO. None of the cocktails were significantly lower than Dyngo4a alone, and ambroxol/Dyngo4a was almost equivalent to DMSO, indicating that ambroxol negated the inhibitory effects of Dyngo4a (Figure 6A).

**Figure 6.**
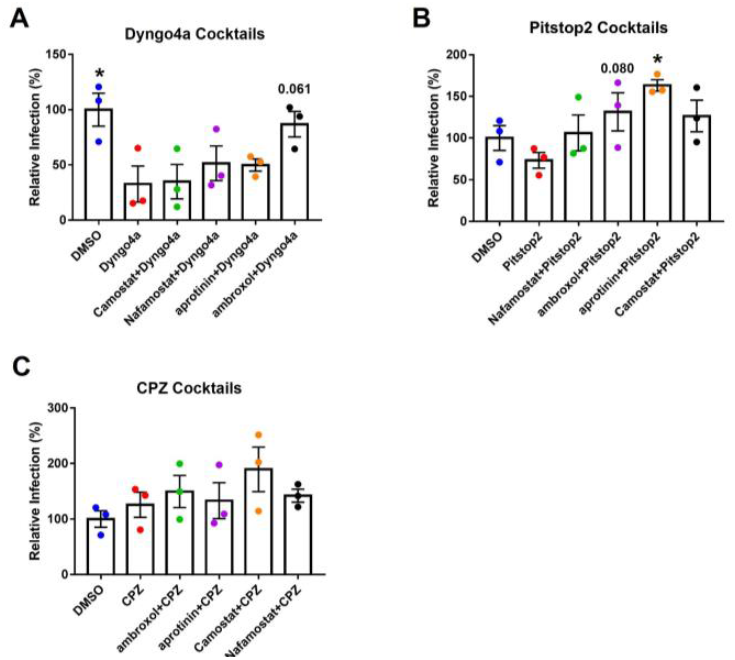
Effects of targeting both TMPRSS2 and endocytosis on infection. Graphs show data normalized to the DMSO control group. TMPRSS2 inhibitors were paired with (A) Dyngo4a, (B) Pitstop2, or (C) CPZ to determine if mixtures of TMPRSS2 and endocytosis inhibitors were effective in blocking infection. In post-hoc testing, group means were compared to the Dyngo4a group. Dyngo4a alone significantly lowered infection relative to the DMSO control. The addition of aprotinin to pitstop2 significantly increased infection. CPZ mixtures were not significantly different than the DMSO control. Data are the means ± SEM of three independent experiments. *p < 0.05. One-way ANOVAs were performed on raw data.

Pitstop2 alone lowered infection when compared to the DMSO control, while the addition of TMPRSS2 inhibitors increased infection above that observed with Pitstop2 (Figure 6B). The addition of aprotinin significantly increased infection, and the increase in ambroxol/Pitstop2 was close to significant (p = 0.08).

CPZ alone did not inhibit SARS-CoV-2 pseudoparticle infection (Figure 6C). None of the cocktail groups showed a significant decrease in infection levels relative to CPZ alone. CPZ cocktails containing TMPRSS2 inhibitors were either similar to CPZ alone or showed increased infection above the DMSO control.

Caveolae-mediated endocytosis, another dynamin-dependent pathway, was targeted using cholesterol inhibitors, which included methyl-β-cyclodextrin (mβCD), nystatin, filipin, and genistein (Figure 7A). Nystatin significantly increased infection compared to the DMSO control. Other cholesterol inhibitors lowered infection by 3-20%, but none were significantly different from the DMSO control.

**Figure 7.**
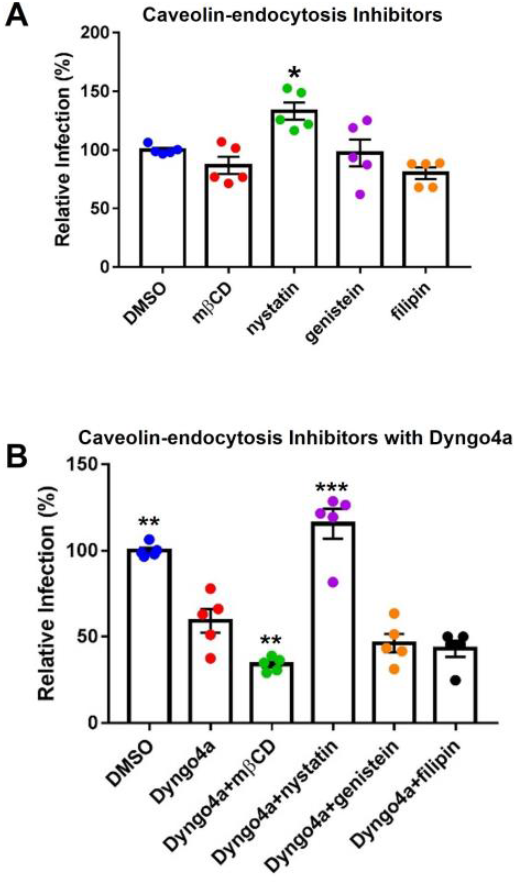
Effects of cholesterol inhibitors on infection. (A) Cholesterol inhibitors did not significantly affect caveolin-mediated endocytosis-based infection, except for nystatin which increased infection. (B) In post-hoc testing, group means were compared to the Dyngo4a group. The addition of mβCD to Dyngo4a significantly lowered infection than individual treatment. Graphs show data normalized to the DMSO control group. Data are the means ± SEM of three independent experiments. *p < 0.05, **p < 0.01, **p < 0.001. Asterisks above the groups indicate comparison with the DMSO control. One-way ANOVAs were performed on raw data.

Cholesterol inhibitors were also mixed with Dyngo4a to block dynamin and caveolin simultaneously (Figure 7B). In Dunnett’s post-hoc test, Dyngo4a was used as the comparative group. Infection in the Dyngo4a group was significantly lower than the DMSO control. Cocktails containing mβCD, genistein, and filipin all lowered infection relative to Dyngo4a; however, only Dyngo4a/ mβCD was significantly lower than Dyngo4a. In contrast, the addition of nystatin to Dyngo4a caused a significant increase in infection relative to Dyngo4a alone.

## 4 Discussion

Despite the documented transmission of SARS-CoV-2 to at least 23 non-human species and the emergence of variants of concern, zoonosis remains an understudied aspect of COVID-19 research (Madhusoodanan 2022). Our data showed that SARS-CoV-2 pseudoparticles readily infected Mv1Lu cells and provide an efficient and safe method for studying the early stages of infection in mink. Combining Mv1Lu cells with SARS-CoV-2 pseudoparticle infection enabled the rapid and efficient screening of potentially therapeutic drugs. Infection was successfully mitigated by endocytosis inhibitors, such as Dyngo4a, but was not affected by TMPRSS2 inhibitors, supporting the conclusion that SARS-CoV-2 virus entered Mv1Lu cells mainly by endocytosis. The effective small molecule inhibitors identified in this study could have therapeutic potential and may be useful in preventing SARS-CoV-2 infection of mink.

### 4.1 Mv1Lu cells as a model for studying SARS-CoV-2 infection

SARS-CoV-2 infection has devastated mink farms worldwide (Molenaar et al. 2020; Cahan 2020), and the infection of mink has many similarities to SARS-CoV-2 infection of humans (Devaux et al. 2021). As examples, the symptoms displayed in mink are similar to those in humans, and the virus is rapidly transmitted in both mink and humans. Moreover, both spillover and spillback events have been reported in mink (Devaux et al. 2021).

Given the importance of mink as a carrier of SARS-CoV-2, it is desirable to have a mink cell line that allows *in vitro* experimentation and evaluation of potentially therapeutic drugs. As originally shown by Heller et al. (2006), Mv1Lu cells from mink express the ACE2 receptor. We extended this observation by showing that Mv1Lu cells are a valuable *in vitro* model for studying SARS-CoV-2 infection in mink, characterizing the SARS-CoV-2 entry mechanisms in these cells, and screening drugs that could inhibit infection. ACE2 was robustly expressed in Mv1Lu cells at both the RNA and protein levels, TMPRSS2 protease was both expressed and enzymatically active, and SARS-CoV-2 pseudoparticles were efficiently taken up by endocytosis. Mv1Lu cells, which were originally isolated from fetal lungs and spontaneously immortalized, are easy to culture and readily available to any lab (Henderson et al. 1974). Mv1Lu cells allow convenient and rapid screening of drugs that can potentially prevent SARS-CoV-2 infection in mink. SARS-CoV-2 pseudoparticles, which cannot replicate, can be generated in a Biosafety Lab 2 cabinet and provide a safer alternative than using live virus. This infection system has been adapted to other cell types and species (Song et al. 2023). Pseudotyping has been used previously to study virus-host receptor binding (Lu et al. 2014; Wang et al. 2008) and can be applied to Mv1Lu cells, which are a robust *in vitro* model for studying coronavirus infection in mink.

### 4.2 Small molecule inhibitor screening showed that endocytosis is the major viral entry pathway in Mv1Lu cells

The use of small molecule inhibitors with Mv1Lu cells enabled characterization of the viral entry pathway and identification of potential therapeutics for preventing infection (Figure 8). In humans, small molecule inhibitors have been used previously to show that SARS-CoV-2 enters host cells by both the TMPRSS2-based fusion and endocytosis pathways (Zhao et al. 2022; Peacock et al. 2022; Neerukonda et al. 2022). Our study showed that in mink some small molecule inhibitors were effective (Dyngo4a, mβCD), some were not (ambroxol, Camostat, Nafamostat, aprotinin), and some worsened infection (CPZ, nystatin), demonstrating the importance of testing drugs before using them therapeutically (Figure 8). Dyngo4a, which blocks the dynamin GTPase involved in various endocytic pathways (Macia et al. 2006), was the most effective small molecule inhibitor. Pitstop2, mβCD, and filipin, which block endocytosis (von Kleist et al. 2011; Vercauteren et al. 2010), were also effective in preventing pseudoparticle infection of Mv1Lu cells. Pitstop2 targets the clathrin heavy chain and prevents it from interacting with adaptor proteins required for clathrin-coated pit formation (Alkafaas et al. 2022). Other inhibitors blocked caveolae-mediated endocytosis by various mechanisms. For instance, mβCD inhibits endocytosis by depleting cholesterol (Rodal et al. 1999; Rejman et al. 2005; Zidovetzki and Levitan 2007), which leads to a decrease in the number of caveolae in other cell types (Dreja et al 2002). Although the mechanism of action of nystatin is similar to mβCD, it was cytotoxic in Mv1Lu cells, indicating it is not suitable for use in mink. Filipin interacts with membrane cholesterol and prevents the close packing of acyl chains (Alkafaas et al. 2022), and therefore, was effective in decreasing infection in mink. Genistein, an FDA-approved pharmacological inhibitor of Src kinase-dependent phosphorylation of caveolin-1, (Zhang et al. 2018), decreased infection in mink. Our results show that most endocytosis inhibitors were effective in preventing SARS-CoV-2 pseudoparticle infection in Mv1Lu cells; however, they varied in their efficacy. The results with nystatin demonstrate the importance of drug screening, as not all endocytosis inhibitors can be assumed to be effective. Overall, our data support the conclusion that the SARS-CoV-2 virus enters mink Mv1Lu cells mainly via dynamin-dependent endocytosis, which includes both clathrin-dependent and -independent pathways (Rennick et al. 2021).

**Figure 8.**
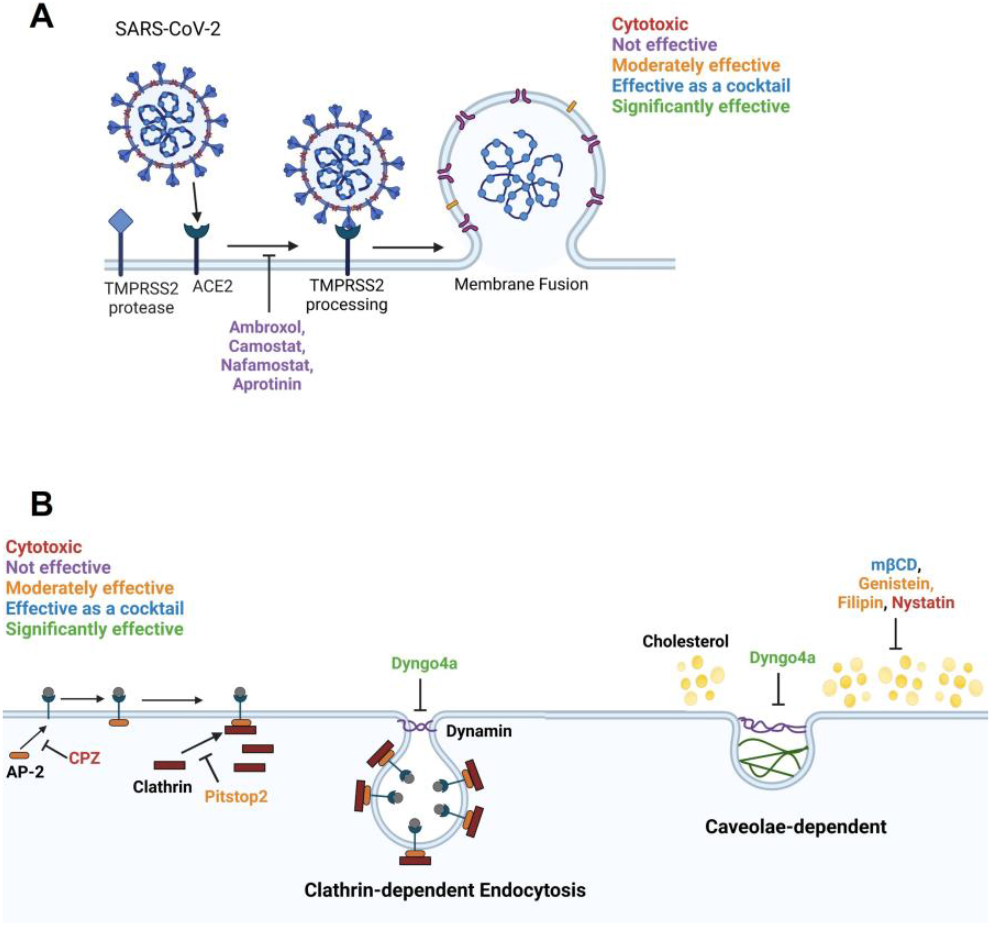
Diagram summarizing the effects of inhibitors on SARS-CoV-2 entry into Mv1Lu cells. (A) The membrane fusion entry pathway. None of the inhibitors effectively blocked this pathway. (B) The dynamin-dependent endocytosis pathway. The clathrin-mediated endocytosis pathway can be blocked by targeting various intermediates. Pitstop2, which inhibits clathrin from binding to AP-2, was moderately effective and Dyngo4a, which reduces dynamin GTPase activity, was significantly inhibitory. In contrast, CPZ, which blocks binding of AP-2 to the ACE2 receptor, was cytotoxic. Caveolae-dependent endocytosis can be blocked by targeting cholesterol. mβCD, filipin, and genistein were moderately effective. mβCD, which depletes cholesterol from the plasma membrane, was significantly effective as a cocktail with Dyngo4a. Filipin loosens the packing of acyl chains by interacting with cholesterol. Genistein blocks the phosphorylation of caveolin. Nystatin functions similarly to mβCD but was cytotoxic to Mv1Lu cells.

CPZ, a FDA-approved drug for treating psychiatric disorders (Daniel et al. 2015; Wu et al. 2016), blocks clathrin-coated vesicle formation; however, it was not effective in inhibiting SARS-CoV-2 pseudoparticle infection and showed cytotoxicity in Mv1Lu cells. CPZ was effective in blocking live virus infection in human A549 cells (Weston et al. 2020; Plaze et al. 2021). Since psychiatric patients receiving CPZ were observed to show a lower prevalence of COVID-19 (Plaze et al. 2020), clinical trials are registered in Egypt and France to determine if administration of CPZ decreases the likelihood of contracting COVID-19 in humans (Stip et al. 2020). Our CPZ results suggest that species can differ in their sensitivity to this drug and that efficacy varies with species.

SARS-CoV-2 can also enter host cells by an alternate pathway that involves viral spike protein binding to the ACE2 receptor and subsequent cleavage of spike by the TMPRSS2 protease (Hoffman et al. 2020). Most prior zoonotic studies on this pathway have focused on ACE2 conservation and ACE2 affinity to predict the host range of SARS-CoV-2 (Damas et al. 2020; Sun et al. 2021). More recent SARS-CoV-2 studies have also demonstrated the importance of testing TMPRSS2 activity as well as expression (Shrimp et al. 2020; Phandthong et al. 2022; Phandthong et al. 2023). In our study, TMPRSS2 inhibitors did block enzymatic activity, but they did not reduce SARS-CoV-2 infection, supporting the conclusion that this pathway was not important in Mv1Lu cells. However, Camostat and Nafamostat did reduce SARS-CoV-2 infection in human Calu-3 cells (Hoffman et al. 2020; Wang et al 2020), again demonstrating the importance of testing inhibitors on the species of interest. Together, our viral pseudoparticle infection assay and small molecule inhibitor data showed that Mv1Lu cells used endocytosis for SARS-CoV-2 entry, but not the TMPRSS2-based fusion pathway (Figure 8).

Our study provides a framework for future work on viral entry mechanisms and development of therapeutics for SARS-CoV-2 infections in zoonotic species. While our study focused on the wild-type spike strain (Wuhan-Hu-1), our protocol could be adapted for use with other variants and tested using other zoonotic species (Song et al. 2023). Finally, our approach could be used to explore an alternative endocytic entry pathway facilitated by soluble ACE2 (Yeung et al. 2021), highlighting the versatility of this infection system for drug screening.

### 4.3 Testing the cocktail effects is important to carefully evaluate toxicity and efficacy

Drug cocktails that target multiple viral proteins or multiple steps in the viral pathway are a promising type of therapeutics for treating viral diseases. Drug combinations can be more effective than single drugs in blocking viral entry by eliminating drug-resistant variants (Yang and Rao 2021). It is possible that when one entry pathway is blocked, another pathway may be activated. Computational studies on SARS-CoV-2 infection predicted that targeting both TMPRSS2 and cathepsin lowered the infectability more than targeting either pathway alone (Padmanabhan et al. 2020). Similarly, in our study, infection was lowered more when Dyngo4a and mβCD were combined than when either was used independently. Dyngo4a (Rennick et al. 2021) and mβCD (Rodal et al. 1999) were reported to block both clathrin and caveolin-mediated endocytosis, which appeared to be the case in our study. Adding mβCD may have acted by altering the fluidity of the plasma membrane, as shown by others (Zidovetzki and Levitan 2007; Dreja et al. 2002).

### 4.4 Administration and delivery of validated small molecule inhibitors

Our study supports the conclusion that targeting dynamin and cholesterol with small molecule inhibitors, Dyngo4a and mβCD, can offer a new approach for the prevention of SARS-CoV-2 infection in mink and complement COVID-19 vaccines. mβCD is already approved as an additive to food by the Flavor Extract Manufacturers Association (FEMA) and is “generally recognized as safe” (FEMA GRAS 4028). mβCD is also currently used to increase the solubility of non-polar drugs for various routes of administration in humans (Uekama et al. 1998). Dyngo4a is not yet FDA-approved, but many preclinical reports propose Dyngo4a as a candidate therapeutic. For example, in an *ex vivo* perfused mouse heart model of ischemia/reperfusion injury, Dyngo4a inhibited mitochondrial fission by targeting mitochondrial dynamin (Drp1), leading to increased cardiomyocyte survival (Gao et al. 2013). Moreover, in the corneal epithelium (human *in vitro* and mouse *ex vivo*), oxidative stress and hyperosmolar stress (in the ER) can be alleviated by Dyngo4a (Webster et al. 2018; Martinez-Carrasco et al. 2020; Martinez-Carrasco and Fini 2023). In our study, at Dyngo4a concentrations that were effective in blocking SARS-CoV-2 pseudoparticle entry, other cellular processes did not appear to be affected. For example, Mv1Lu cells continued to divide and appeared morphologically normal in the presence of Dyngo4a.

Nasal administration with nebulizers is one promising approach (Eedara et al. 2021) to effectively deliver therapeutic drugs to mink. The delivery of Dyngo4a and mβCD via nasal spray directly introduces the drugs to the desired target (respiratory system) in contrast to drugs that are injected or ingested. Given the ability of mβCD to solubilize various drugs, it is currently widely used as an excipient for enhancing drug delivery via nebulization (Evrard et al. 2004; Chandrama Singh et al. 2022). Nebulization with Dyngo4a and mβCD could also be used in conjunction with other therapies, such as antisense oligos, which could be delivered simultaneously or in tandem (Zhu et al. 2022). Our results indicate that targeting the early infection steps of SARS-CoV-2 virus holds promise for the development of therapeutics that are effective in treating mink populations, a strategy that has been suggested for other animals by other virologists (Mercer et al. 2010; Mazzon and Marsh 2019). Our data further show that when repurposing drugs to prevent SARS-CoV-2 infection in mink, focus should be on the endocytosis pathways.

### 4.5 Conclusions and Significance

Through vaccinations and other public health measures, the COVID-19 pandemic has been brought under reasonable control for the human population. However, the virus is harbored by many non-human animals, where it can mutate into dangerous variants that could spill back into humans. Understanding SARS-CoV-2 infection of mink is important to prevent future spillback of mutated virus from mink to humans. The problem of zoonotic transmission needs to be addressed by: (1) improving monitoring and testing of non-human species, (2) implementing measures to prevent zoonotic transmission from animals to humans, and (3) developing vaccines and/or other treatments that can be used to protect both human and animal populations. Our study addressed the third point, and it is the first to report that SARS-CoV-2 enters a non-human host using the endocytic pathway and that targeting dynamin with Dyngo4a can prevent entry. Though animal vaccines have been developed to facilitate the management of COVID-19 (Hoyte et al. 2022), our study suggests that the blockage of entry is another potentially powerful strategy. In addition, evaluating the toxicity and efficacy of individual or combined drugs is essential because mixtures may be more or less effective. Effective inhibitors identified in this study could be delivered in a nebulized form, which should be feasible with mink and relatively inexpensive. Our results provide a foundation for developing novel therapeutics to prevent SARS-CoV-2 entry into mink, possibly other animals, including humans, and may inform public health policy. The data from our mink study could reduce the risk of continued transmission between animals and humans and prevent the development of more dangerous variants.

## 5 Additional Requirements

None

## 7 Author Contributions

AS and PT contributed to conception, data interpretation, manuscript writing, and revision of the submitted version. PT was responsible for project administration and funding acquisition. AS and RP contributed the 293T sample preparation, data collection, and processing. AS was responsible for Mv1Lu sample preparation, data collection, and processing.

## 9 Acknowledgments

Figure 1 and 8 were created with Biorender.com. We thank Claudia Osuna and Jack Ona for making some of the batches of viral pseudoparticles. We thank the UCR Stem Cell Core for providing access to the Novocyte Flow cytometer.

## 11 Supplementary Material

Uploaded as a separate file.

## 12 Data Availability Statement

The original contributions presented in the study are included in the article, further inquiries can be directed to the corresponding author.

