## Supplementary Figure 1 for "Endocytosis Inhibitors Block SARS-CoV-2 Pseudoparticle Infection of Mink Lung Epithelium"

SARS-CoV-2 Pseudoparticles Enter the Mink Lung Epithelium via Endocytosis

### 1 Supplementary Figure

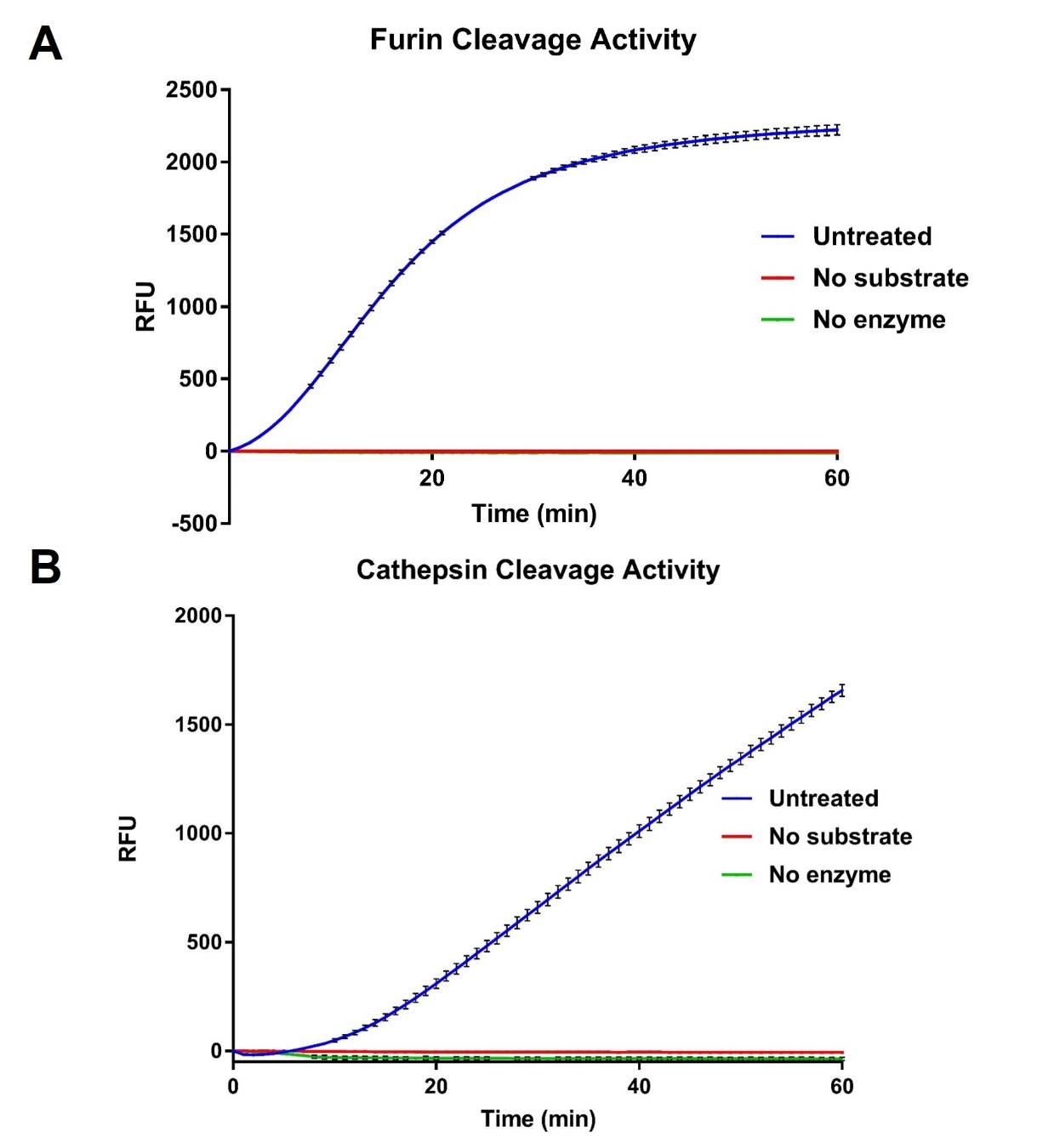

**Supplementary Figure 1.** Mink Mv1Lu cells have robust protease activities that are essential to SARS-CoV-2 infection. (A) furin and (B) cathepsin L cleavage activities were assessed using a fluorogenic substrate. Both furin and cathepsin L activities were high in Mv1Lu cells. DMSO, solvent control; RFU, relative fluorescence unit or normalized fluorescence intensity. Each line is the mean of three independent experiments. Error bars represent SEM.
